# Telomere shortening produces an inflammatory environment that promotes tumor invasiveness in zebrafish

**DOI:** 10.1101/844084

**Authors:** Kirsten Lex, Mariana Maia Gil, Margarida Figueira, Marta Marzullo, Bruno Lopes-Bastos, Kety Giannetti, Tania Carvalho, Miguel Godinho Ferreira

**Author notes:** These authors contribute equally in this work (Kirsten Lex, Mariana Maia Gil).

## Abstract

Cancer incidence increases exponentially with age, when human telomeres are shorter. Similarly, telomerase mutant zebrafish (*tert*) have premature short telomeres and anticipate cancer incidence to younger ages. However, because short telomeres constitute a road block to cell proliferation, telomere shortening is currently viewed as a tumor suppressor mechanism and should protect from cancer. This conundrum is not fully understood. In our current study, we report that telomere shortening promotes cancer in a non-cell autonomous manner. Using zebrafish chimeras, we show increased incidence of invasive melanoma when WT tumors are generated in *tert* mutant zebrafish. *tert* zebrafish show increased levels of senescence (*cdkn2a* and *ink4a/b*) and inflammation (*TNF-*α). In addition, we transferred second generation *tert* blastula cells into WT to produce embryo chimeras. Cells with very short telomeres induced senescence and increased neutrophil numbers in surrounding larval tissues in a non-cell autonomous manner, creating an inflammatory environment. Considering that inflammation is pro-tumorigenic, we transplanted melanoma-derived cells into second generation *tert* zebrafish embryos and observed that tissue environment with short telomeres leads to increased micrometastasis. To test if inflammation was necessary for this effect, we treated melanoma transplants with non-steroid anti-inflammatory drugs and show that higher melanoma invasiveness can be averted. Thus, apart from the cell autonomous role of short telomeres in contributing to genome instability, we propose that telomere shortening with age causes systemic chronic inflammation leading to increased tumor incidence.

**Significance Statement:** Cancer incidence increases exponentially in human midlife. Even though mutation accumulation in somatic tissues results in increased tumorigenesis, it is currently not understood how aging contributes to cancer. Telomeres, the ends of eukaryotic linear chromosomes, shorten with each cell division. Here we show that telomere shortening contributes to cancer in a non-cell autonomous manner. Using embryo chimeras of telomerase deficient zebrafish generated from melanoma-prone fish, we show that tumors arise more frequently and become more invasive in animals with shorter telomeres. Telomere shortening gives rise to increased senescence and systemic inflammation. We observed increased melanoma metastasis dissemination in zebrafish larvae with very short telomeres. Thus, telomere shortening similar to human aging, generates a chronic inflammatory environment that increases cancer incidence.

## Introduction

Cancer incidence increases exponentially in the mid-decades of human life (1). Although mutations are required to build-up during tumorigenesis, the overall post-reproductive incidence opens the possibility of organism-based causes for the increase of cancer with age. Due to absence of telomerase expression in most somatic tissues, telomeres shorten as we grow older (2). Telomeres constitute the ends of eukaryotic chromosomes and are constituted by repetitive DNA sequences (TTAGGG)_n_ recognized by a protein complex called shelterin (3). This structure prevents chromosome-ends from being recognized as deleterious DNA double strand breaks while counteracting their slow attrition, resulting from the “end-replication problem” by recruiting telomerase. Humans are born with telomeres between 10-15 kb long (4) and, due to continuous cell divisions, telomeres may reach a critical length. As cell division reaches the Hayflick limit, telomeres are recognized as DNA damage and block cell proliferation either by undergoing senescence or apoptosis (5–7). Since short telomeres block cell division, telomere shortening is considered as a tumor suppressor mechanism by preventing excessive cell proliferation. Indeed, telomerase is frequently re-activated in the majority of cancer cells, allowing for cell immortalization thereby escaping replicative senescence. In line with this idea, anti-telomerase therapies are currently undergoing clinical trials for cancer therapy (8).

Countering the tumor suppressor hypothesis, telomere shortening may lead to genome instability, a hallmark of cancer. Because loss of telomere protection results in breakage-fusion-bridge cycles, the ensuing genome instability may contribute for age-dependent tumorigenesis (9). An extreme example of the pro-tumorigenic effect of short telomeres occurs in “telomeropathies”. People carrying mutations in telomerase or related proteins have pathologically short telomeres in early life (10, 11). Despite exhibiting pathologies related to deficiencies in cell proliferation, patients also suffer from an increased cancer risk (12). Similarly, our work on the telomerase mutant zebrafish, which undergoes premature telomere shortening, revealed that they anticipate cancer incidence to early life (13). Even though short telomeres positively correlate with increased tumorigenesis in both humans and zebrafish, it is not yet understood how telomere shortening may lead to cancer.

Telomere shortening has consequences beyond the cellular level. As cells approach replicative senescence, DNA damage emanating by short telomeres initiate a cascade of events that expands to the extracellular environment. Senescent cells were shown to release a set of molecules termed senescence-associated secretory phenotype (SASP) (14). SASP was described *in vitro* and is mainly constituted by chemokines, growth factors, extra cellular matrix remodelers and other inflammatory factors, capable of modulating cell environment. These molecules were posteriorly shown to influence the ability of other cells to divide, potentially having a pro-tumorigenic effect (15). Consistently, repeated wounding in zebrafish stimulates inflammatory responses, which were shown to promote cancer progression (16, 17). Therefore, we hypothesize that telomere shortening contribution to tumorigenesis may have a non-cell autonomous component. In aging organisms, cells undergoing replicative senescence would comprise a source of SASP/inflammatory factors creating a pro-tumorigenic environment. In agreement with our hypothesis, population studies have associated the long-term use of anti-inflammatory agents (acetylsalicylic acid) and a reduction risk of several cancers (18–20).

Here we show that tissues containing cells with short telomeres promote increased cancer incidence in a non-cell autonomous manner. Using chimeric zebrafish, we observed that telomerase-proficient melanocytes expressing HRAS give rise to more melanoma tumors when surrounded by *tert* mutant cells. Melanomas developed in this environment exhibited high invasiveness as observed by histopathology. In agreement, using zebrafish tumor transplants, we show that HRAS melanoma cells expand faster when injected into second-generation (G2) *tert* mutant larvae. Both adult G1 *tert* and G2 *tert* larvae have higher levels of senescence and SASP/inflammation. G2 *tert* cells injected into WT embryos stimulate senescence and inflammation in a non-cell autonomous manner. Chemical inhibition of inflammation in G2 *tert* embryos rescued the invasiveness capacity of melanoma cells. Thus, cells with short telomeres are capable of inducing senescence and inflammation, creating a pro-tumorigenic environment that results in higher cancer invasiveness.

## Results

### *tert* mutant environment causes higher tumor incidence in a non-cell autonomous manner

Similar to mammals, zebrafish tumor microenvironment (TME) modulates cancer behavior (21, 22). Tumors may be inhibited or enhanced as a consequence of the dynamic crosstalk between cancer and surrounding cells. We, therefore, asked what were the effects of a TME with short telomeres on emergent tumors.

In order to study the non-cell autonomous effects of TME telomere shortening in cancer, we wanted to separate telomerase expression of pre-cancer cells from their surrounding tissues and, for this purpose, we generated chimeric zebrafish using early-developmental embryo transplants. We used a melanoma zebrafish model (*mitfa*:HRAS) developed by the Hurlstone lab that exhibits full penetrance by 3 months of age (23). We chose this model since it did not require an initial *tp53* dysfunction to form tumors and we had previously shown that loss of p53 function rescues *tert* zebrafish mutants (24). Blastula cells from donor embryos capable of giving rise to melanoma were transplanted into WT or *tert−/−* recipients (Fig 1A). In addition, recipient embryos had a *casper* genetic background (*mitfa*^*w2/w2*^; *mpv17*^*a9/a9*^), and lacked the ability to produce melanocytes. Consequently, all melanoma could only arise from donor cells. Embryo chimeras then were allowed to grow into adulthood and studied for tumor incidence. As expected, we observed the development of melanoma lesions, typically in the anal fin region of both WT and *tert−/−* recipient fish (Fig. 1B). However, by 30 weeks, a time when *tert−/−* associated lethality is still low (<20%), 20% of WT chimeras developed tumors, while ca. 50% of *tert−/−* chimeras exhibited melanoma (Fig. 1C, p<0.05). Thus, we found that *tert−/−* recipients significantly increased tumor incidence by ca. 2-fold (Hazard ratio after Mantel-Haenszel calculation: 2.0 when compared with WT fish).

**Figure 1.**
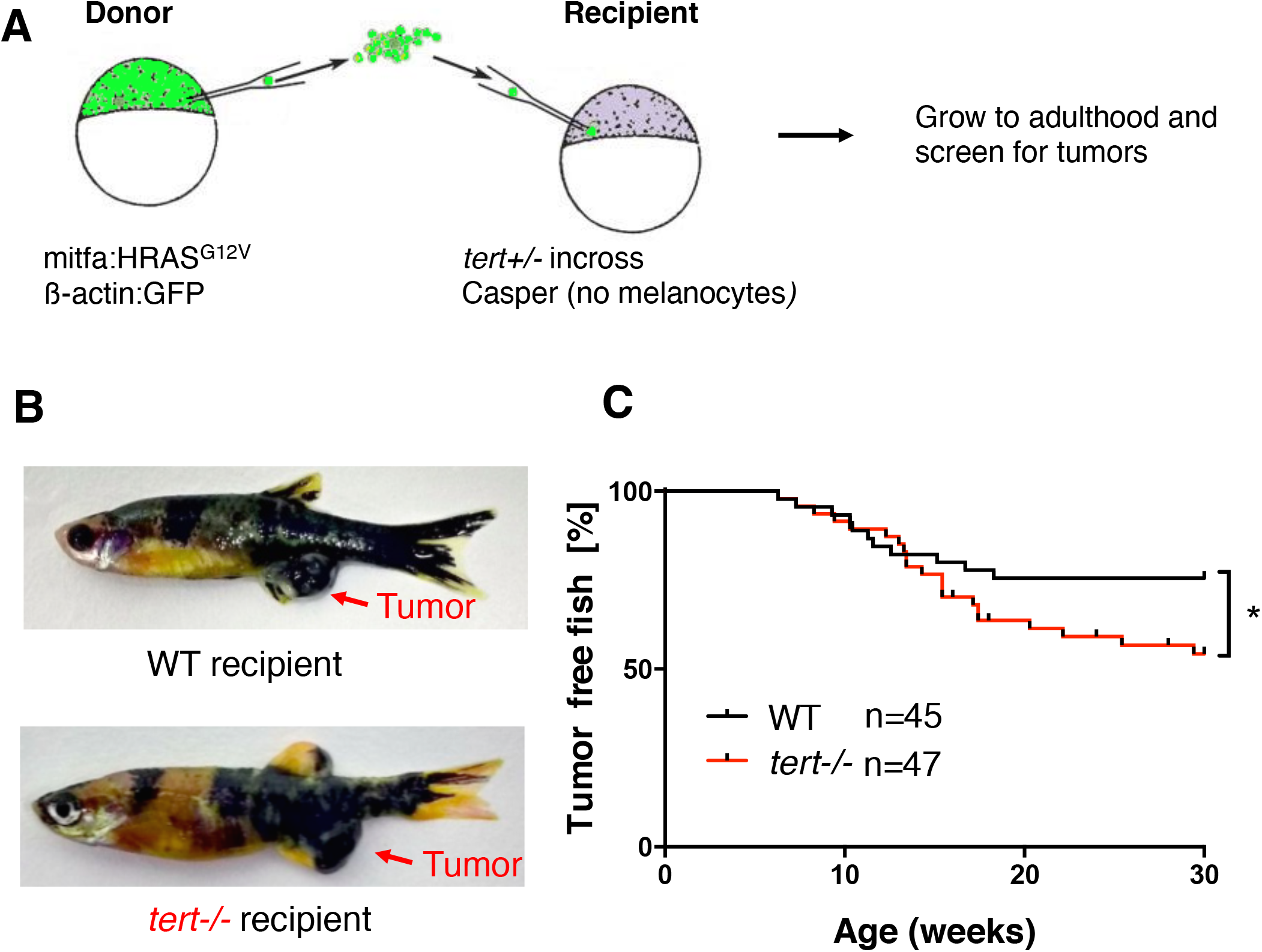
Short telomeres promote tumorigenesis in a non-cell autonomous manner. A) Experimental setup for the generation of zebrafish chimeras. Donor cells are transplanted from a Tg(*mitfa:HRAS*^*G12V*^; β*-actin:GFP*) embryo at the blastula stage into embryos resulting from an incross of *tert+/−*; Casper zebrafish. B) Representative images of adult chimera zebrafish harboring melanoma in either WT or *tert−/−* recipients. C) Melanoma occurrence over time in chimeric fish. *tert−/−* recipient fish have a higher risk of tumorigenesis than WT recipient fish (p<0.05).

A possible explanation for the observed differences of tumor development in a WT vs. *tert−/−* environment is cell competition. Wildtype tumor-prone cells could be fitter and more efficient in outcompeting *tert* mutant recipient cells, possibly due to higher proliferation rates. Thus, fitter donor cells could produce higher number of melanocytes expressing HRAS in *tert* mutant recipients and, subsequently, lead to a higher tumor incidence. To test this hypothesis, we quantified the number of melanocytes at two stages of embryo development at 3- and 11-days post-fertilization (dpf) in both *tert* mutant and WT recipients. Contrary to our hypothesis, we observed no significant increase in the number of melanocytes in *tert−/−* recipients as compared to WT during developmental stages (Supplementary Figure 1A-B). In case growth differences would only be visible at later stages, we quantified the surface area covered by the melanocytic lesions in adult animals. Percentage of pigmentation was quantified for WT and *tert−/−* zebrafish (Supplementary Figure 1C-D). Similar to the results obtained in larvae, although there was variation between individuals, we did not observe significant differences when comparing host genotypes. Together, our data indicates that a *tert* mutant TME increases tumor incidence in a non-cell autonomous manner, suggesting that telomere shortening has a systemic role in cancer beyond the one described in genome stability.

### Tumors progress faster in *tert* mutant TME

Among the hallmarks of cancer, one qualitative difference between cancers relies on the capacity to invade different tissues. Zebrafish chimeras bearing melanoma were analyzed by histopathology and ranked according to their staging and invasiveness. Overall, 84% of samples (N=43) that were macroscopically defined as tumors were confirmed as malignant tumors in histopathological analysis (Fig. 2 A-C). The remaining samples were staged as benign tumors or melanosis. The large majority of tumors in *tert−/−* recipients were invasive (80%, N=10; Fig. 2 B). In comparison, only 22% of tumors exposed to a wildtype environment (N=9) were determined as invasive (Fig. 2 B). A similar result was found when malignant tumors were scored for the presence of cellular atypia. Cellular atypia describes cytologic structural abnormalities and is a marker for more transformed cancers and more advanced staging (25). Whereas 71% of tumors in a *tert−/−* environment (N=7) exhibited moderate levels of cellular atypia (Fig. 2 C), all tumors in WT recipients showed low levels (N=5). These results indicate that melanoma developed in *tert−/−* recipients progress faster, reaching advanced stages faster and becoming more invasive, suggesting that TME telomere shortening not only increase tumor incidence but its progression.

**Figure 2.**
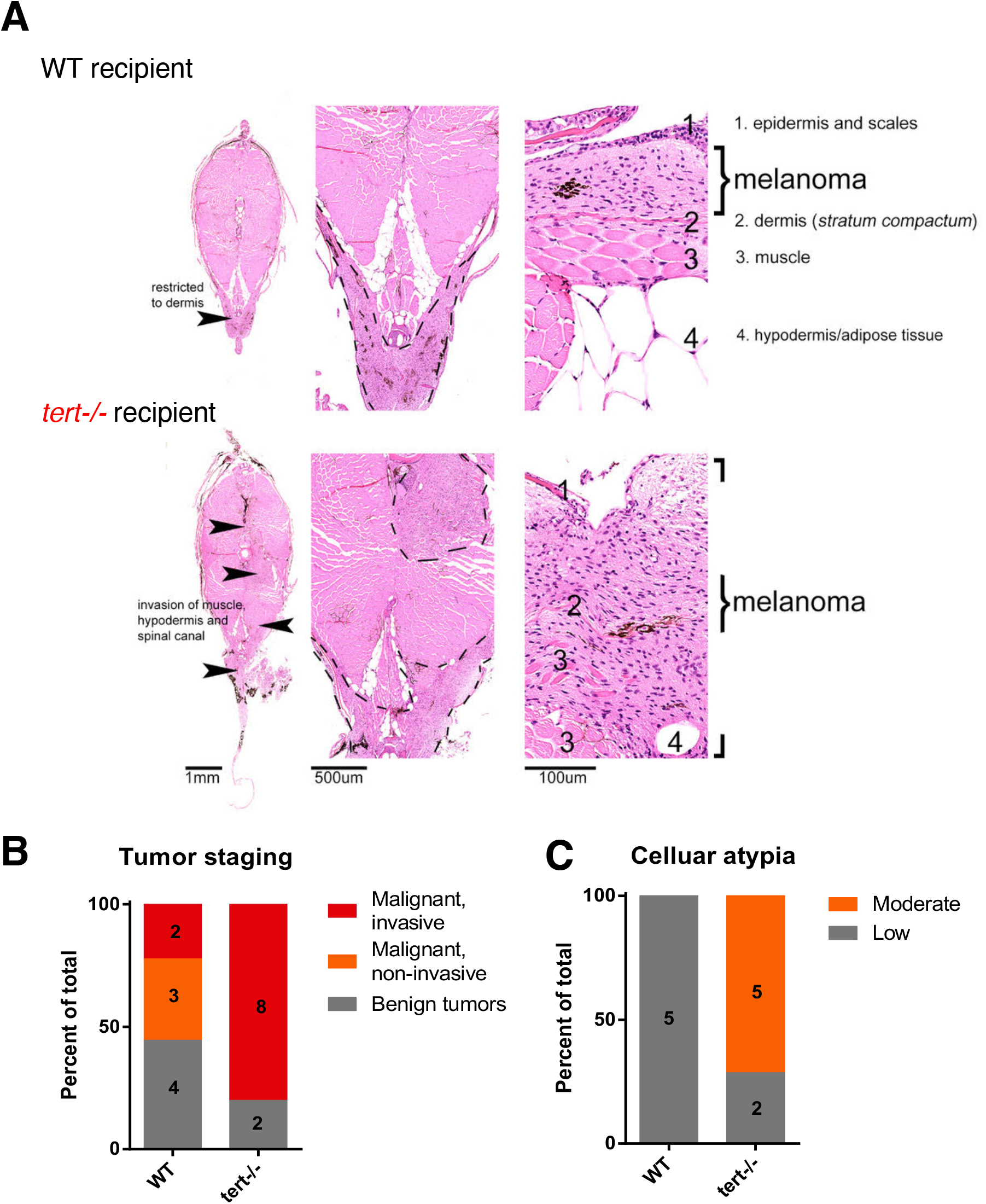
*tert−/−* tissues increase melanoma invasiveness and progression. A) H&E images of melanoma arising in a wildtype (upper panel) or *tert−/−* recipient fish. Strong infiltration into other tissues was typical in *tert−/−* fish but not in wildtype (arrowheads). B) Melanoma were staged according to histopathology into benign lesions (melanosis), non-invasive and invasive malignant tumors. C) Analysis of malignant tumors for cellular atypia. Sample numbers are indicated within the bars.

### Zebrafish melanoma transplants are more invasive in *tert* mutant larvae

We and others have shown that injection of tumor cells in zebrafish larvae constitutes an assay to study invasiveness capacity of cancer cells (26, 27). This constitutes a simpler assay and allows for more expedite manipulations while being amenable to chemical studies.

In order to confirm that *tert−/−* TME promotes tumor invasiveness, we injected melanoma cells derived from HRAS tumors into 2dpf WT and *tert−/−* larvae (Fig. 3A). To ensure that these fish would possess cells with critically short telomeres, we used second generation *tert−/−* (G2 *tert−/−*) resulting from an in-cross of young adult *tert−/−* zebrafish. In contrast to G1 *tert−/−* derived from heterozygous parents, G2 *tert−/−* embryos possess very short telomeres and a high mortality with an average longevity of ~12days (28, 29). We dissected melanomas from HRAS tumors expressing GFP (see Methods) and injected cells into the blood circulation of 2dpf larvae. Injected melanoma cells preferentially accumulate in the tail region from where, depending on their invasiveness capacity, disseminate to neighboring tissues (Fig. 3B). Injected larvae were individually followed over time and the area occupied by GFP cells was quantified (Fig. 3B).

**Figure 3.**
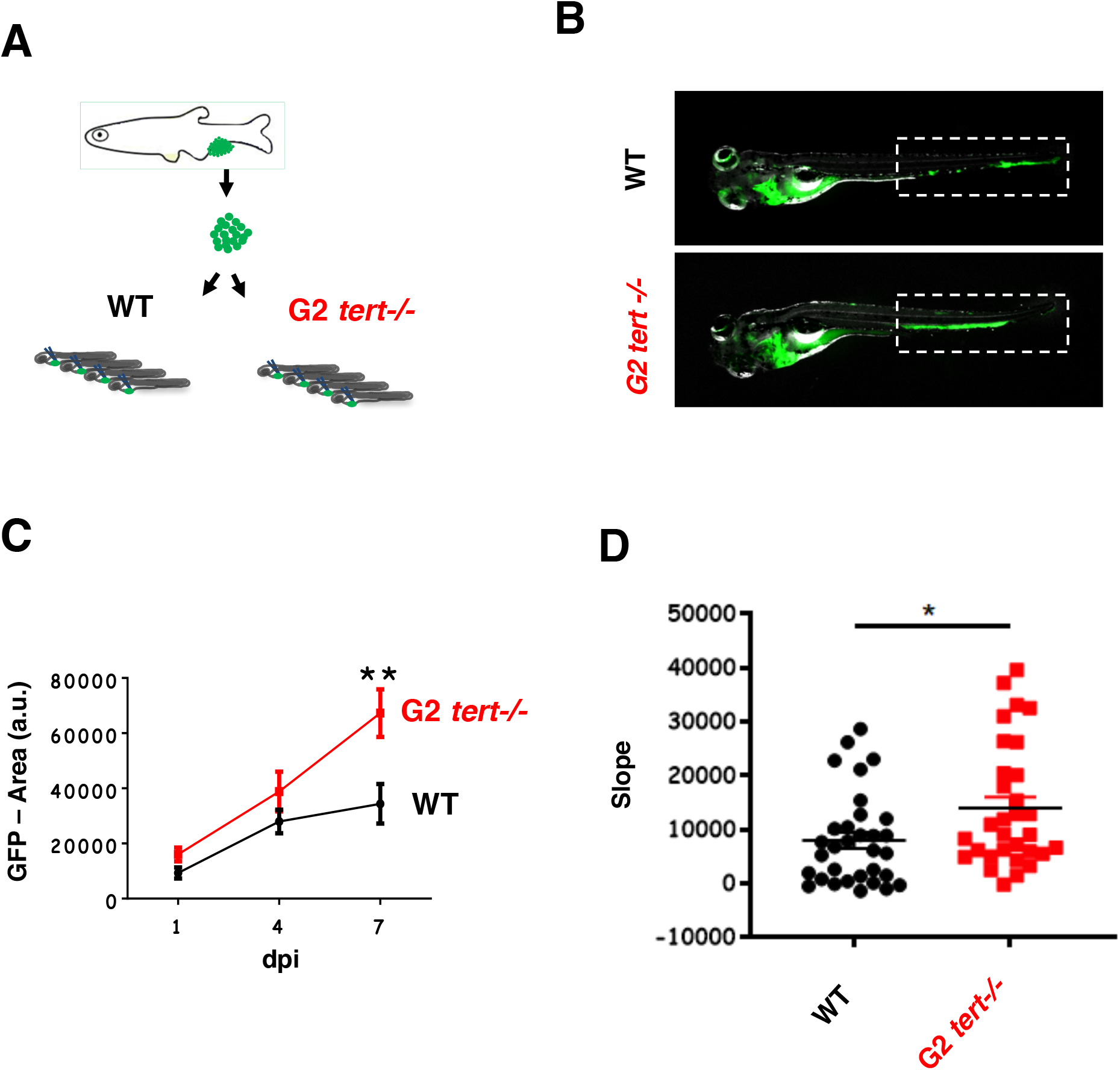
G2 *tert−/−* larvae with very short telomeres exhibit increased melanoma micrometastasis. A) Experimental design for melanoma allotransplants in zebrafish larvae. Melanoma tumors were dissociated from *mitfa:HRAS;* β*-actin:GFP* zebrafish. HRAS melanoma cells were then injected into blood circulation of 2dpf zebrafish larvae. Larvae were kept in embryo medium for 7 days post injection (7dpi). B) Representative images of HRAS melanoma cells spread (green) in WT or G2 *tert−/−* larvae at 7dpi. C) Time-course of HRAS melanoma cells spread in a group of WT and G2 *tert−/−* larvae (p<0.01 at 7dpi, WT N=10 and G2 *tert−/−* N=11). D) Melanoma tumors are more invasive in G2 *tert−/−* larvae (p=0.0205, WT N=32 and G2 *tert−/−* N=31). A linear regression of three time-points (1, 4 and 7 dpi) was used to calculate the slope of melanoma invasiveness. Each dot represents one larvae allotransplant.

If an environment with short telomeres promotes tumor invasiveness, then injected melanoma cells should disseminate more when injected in G2 *tert−/−* when compared to WT larvae. We quantified the GFP-area at 1, 4 and 7 days-post injection (Fig. 3C). We calculated the linear regression between the 3 time-points and obtained a progression slope for the expansion of each grafted melanoma (N=31). We observed that *tert−/−* recipients allowed for a more accentuated progression than the WT ones (Fig. 3D). Thus, our results using tumor transplants indicate that melanoma cells disseminate faster in G2 *tert−/−* than WT larvae, suggesting that telomere shortening in aging individuals could promote tumor progression in a non-cell autonomous manner.

### G2 *tert−/−* cells are senescent and inflammatory and capable of modulating their surrounding environment

Telomere shortening is responsible for replicative cell senescence in human cultured cells (30). Accordingly, we expected that *tert−/−* zebrafish would present increased levels of senescence. Using RT-qPCR for specific genes, we quantified the levels of senescence in *tert−/−* 9month-old adult tissue (intestine) and 4dpf G2 *tert−/−* larvae (whole). As expected, the senescence markers ink4a/b (p15/16) and cdkn1a (p21) levels were significantly higher in both G1 *tert−/−* adults and G2 *tert−/−* larvae than in WT controls (Fig. 4A-B). In addition, using the SA-β-Gal assay, we confirmed higher levels of senescence localized primarily in the head and notochord of G2 *tert−/−* larvae (Fig. 4D).

**Figure 4.**
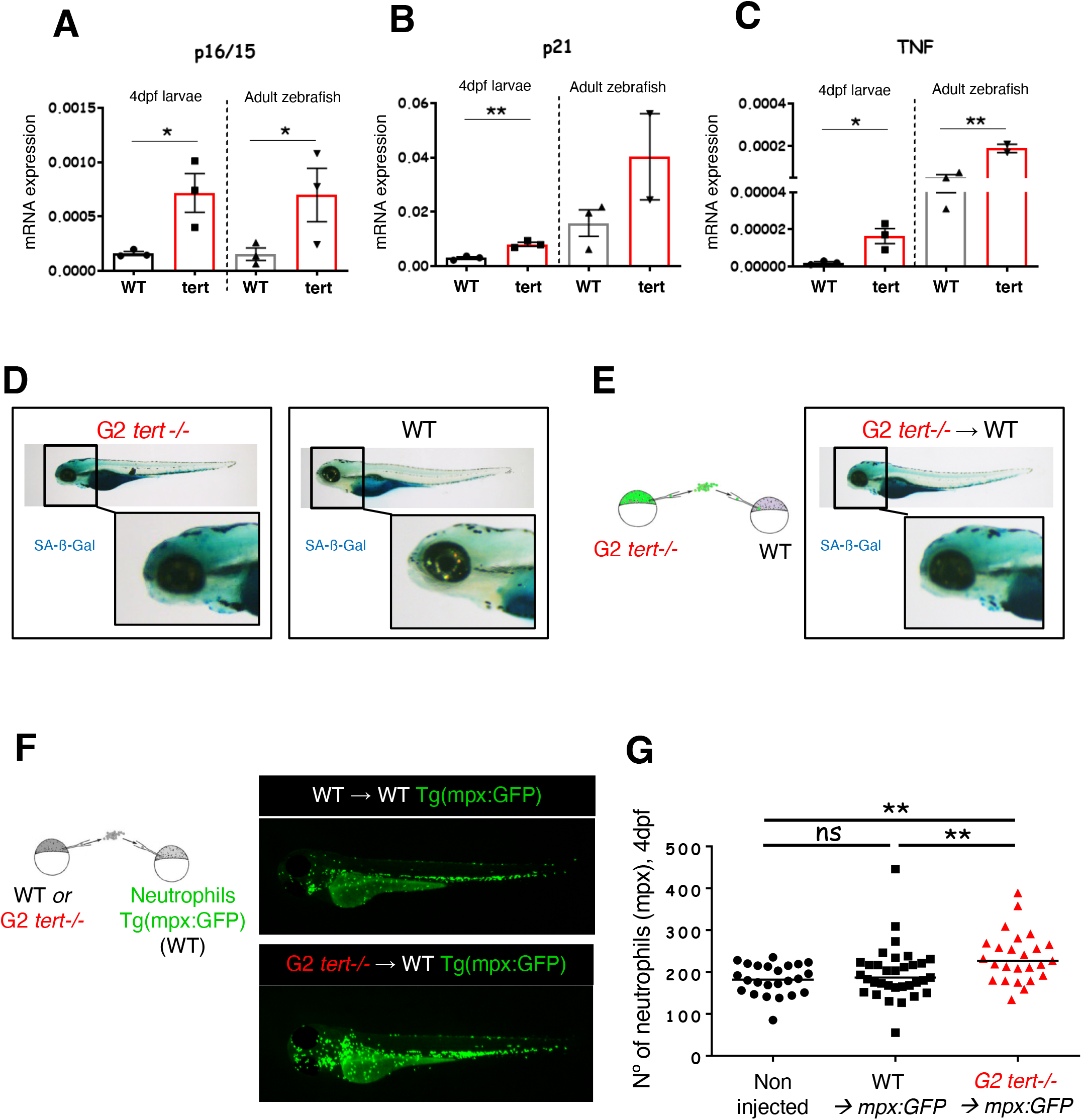
Telomerase deficient tissues present higher levels of senescence and inflammation and modulate their environment. A, B, C) RT-qPCR analysis comparing the expression levels of *ink4ab* (p16/15), *cdkn2a* (p21) and *tnfa* (TNF) of 4dpf WT and G2 *tert−/−* larvae and 9month WT and *tert−/−* adult intestine tissue (* p<0.05, ** p<0.01 N=30). D) Representative images of SA-β-Gal assay comparing WT and G2 *tert−/−* 4dpf zebrafish embryos. Yolk sack staining is nonspecific. E) Scheme for generating chimeras in which G2 *tert−/−* blastula cells are transplanted into WT embryos (G2 *tert−/−*→WT). SA-β-Gal assay showing increased senescence in 4dpf WT embryos with injection of G2 *tert−/−* cells. F) Scheme of G2 chimeras generation where WT or G2 *tert−/−* blastula cells are transplanted into WT Tg(*mpx:GFP*) embryos carrying labelled neutrophils with green fluorescent protein: WT→WT Tg(*mpx:GFP*) vs. G2 *tert−/−*→WT Tg(*mpx:GFP*). Representative images of the chimeras at 4dpf, neutrophils are represented at green; G) Quantification of neutrophils at 4dpf. Non-injected Tg(*mpx:GFP*) were used as controls. Each data point represents one zebrafish (** p<0.01, non-injected N=24, WT N=33 and TERT N=25).

Senescent cells were shown to secrete a set of molecules, known as SASP, mainly composed of inflammatory factors (14, 15). Therefore, we asked if *tert−/−* zebrafish present signs of inflammation. We measured expression levels of TNF-α, one of the main cytokines expressed during an inflammatory response, by RT-qPCR. Indeed, both G1 *tert−/−* adults and G2 *tert−/−* larvae showed elevated levels of TNF-α when compared to WT (Fig. 4C). Interestingly, undisturbed 9month-old WT zebrafish exhibit higher levels of TNF-α than 4day-old larvae, suggesting that aging animals may respond similarly to young *tert−/−* mutants. Together, our results suggest that telomere shortening in zebrafish results in increased senescence and inflammation.

Given the nature of the responses, we wondered if these observations originated from cell-autonomous effects of *tert−/−* cells dispersed through the body or if *tert−/−* cells could modulate their extracellular environment *in vivo* and generate a systemic response. To test if short telomere *tert−/−* cells modulate their extracellular environment, we transplanted GFP-labelled G2 *tert−/−* cells during early-development into WT recipient embryos, thereby generating larvae chimeras (Fig. 4E). Even though we transferred few G2 *tert−/−* cells into developing embryos (<1% as measured by FACS of desegregated embryos at 4dpf), they were sufficient to increase overall SA-β-Gal levels (Fig. 4E). Interestingly, we observed a similar pattern of SA-β-Gal staining in these chimeras as in G2 *tert−/−* larvae (N=22) at the same stage of 4dpf (compare Fig. 4D with 4E). These results suggest that cells derived from G2 *tert−/−* embryos are capable of inducing senescence in a non-cell autonomous manner, thus constituting an example of paracrine SASP.

Since senescent cells secrete pro-inflammatory molecules, we asked if G2 *tert−/−* cells with short telomeres could create an inflammatory environment in newly generated chimeras. To test this, we generated similar embryo chimeras in Tg(*mpx:GFP*) recipient zebrafish that carry GFP-labelled neutrophils (31). As before, we injected both WT and G2 *tert−/−* cells from embryos at blastula stage into Tg(*mpx:GFP*) recipient embryos of the same stage and observed its effects in 4dpf larvae (Fig. 4F, right). Whereas WT cells generated zebrafish larvae (N=33) with similar numbers of neutrophils as un-injected embryos, Tg(*mpx:GFP*) chimeras carrying G2 *tert−/−* cells (N=25) exhibited higher numbers of neutrophils (Fig. 4F, p=0.0075). Thus, since these innate immune cells are key to inflammatory responses, G2 *tert−/−* cells give rise to a systemic inflammatory environment. Together, our results indicate that telomerase deficient zebrafish undergo senescence and produce an inflammatory state. Moreover, we show that this effect is non-cell autonomous with *tert−/−* cells impacting the surrounding tissues modulating their environment, creating a senescent and inflammatory environment.

### Chemical inhibition of inflammation rescues melanoma dissemination in the *G2 tert−/−* mutant larvae

Inflammation can induce transformed cell growth (32). In zebrafish, PGE_2_ produced by innate immune cells via the COX-2 pathway was shown to act as key growth factor at the earliest stages of tumor progression (16, 33). We hypothesized that the inflammatory environment induced by *tert−/−* cells could underlie the enhanced melanoma invasiveness observed in *tert* mutant zebrafish. To test this hypothesis, we treated the previously generated zebrafish melanoma larvae allografts with non-steroid anti-inflammatory drugs (NSAIDs): Aspirin (COX-1 and 2 inhibitor) and Celecoxib (COX-2 specific inhibitor). As previously, we measured melanoma invasiveness by quantifying the GFP area at consecutive timepoints upon melanoma cell injections (1, 4 and 7 dpi). Both WT and G2 *tert−/−* recipients were kept in embryo medium containing Aspirin (30µM) or Celecoxib (25µM) for the duration of the experiment. As previously, we calculated a progression slope of tumor cells per transplanted zebrafish and compared treated vs. untreated larvae (Fig. 5A-B). As previously, control groups showed an increased invasiveness of melanoma cells when transplanted into G2 *tert−/−* (N=31) than in WT larvae (N=32) (Fig. 5D, p= 0.0205). However, upon NSAID treatment, the increased invasion capacity of HRAS cells in G2 *tert−/−* larvae (N=20) decreased to WT levels (N=19) (Fig. 5C-D, Aspirin p=0.7897; Celecoxib: p= 0.1605). Together, our result suggests that the inflammatory environment induced by *tert−/−* cells promotes melanoma invasiveness via the COX-2 pathway. We showed an increase of innate immune cells in larvae containing telomerase deficient cells (Fig. 4F). Thus, in agreement with previous studies (16, 33, 34), we propose that neutrophils, by producing larger amounts of prostaglandins, may enhance melanoma invasiveness.

**Figure 5.**
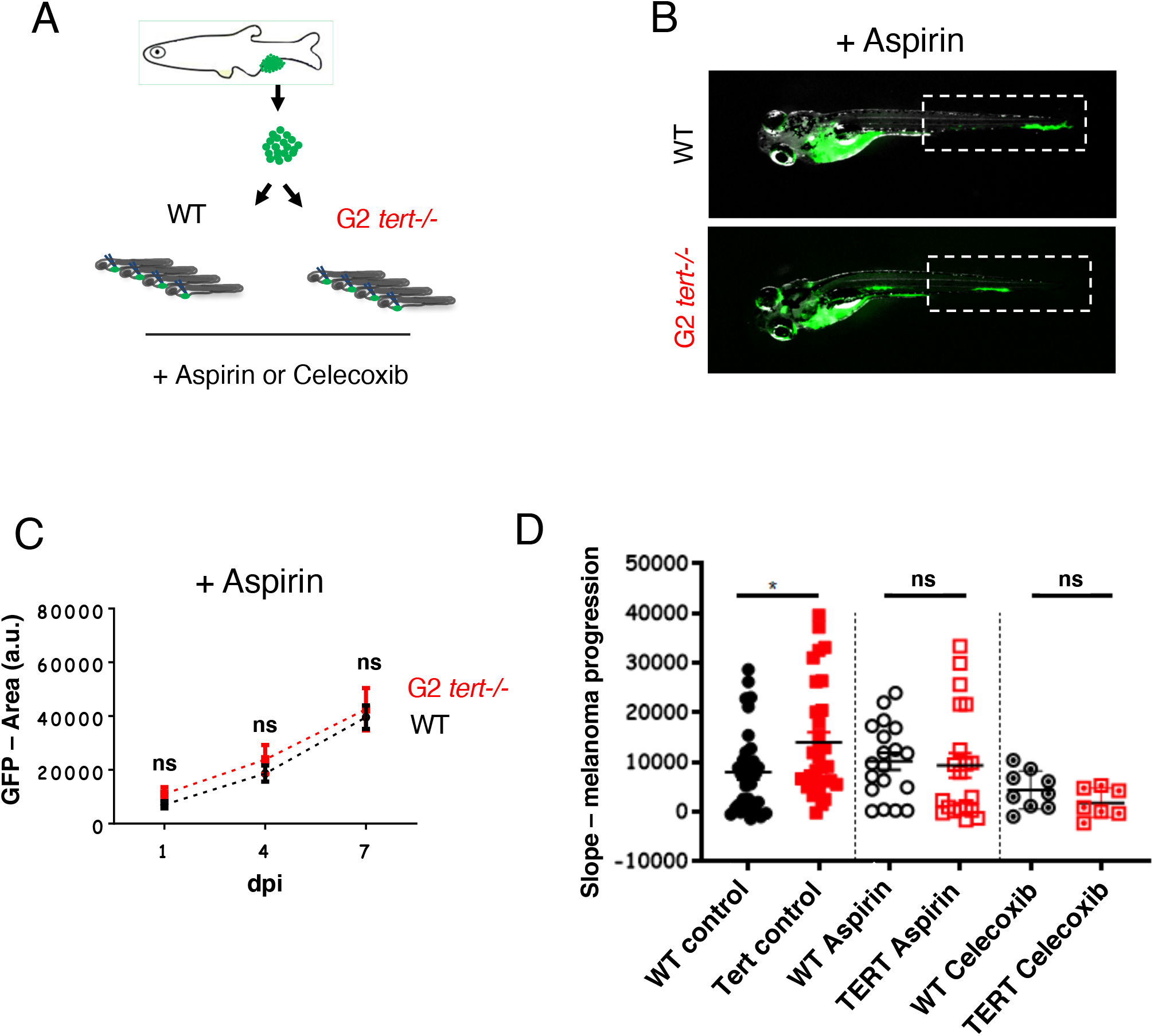
Increased tumor invasiveness in G2 *tert−/−* larvae is rescued by inhibiting inflammation. A) Allotransplants of primary tumor cells extracted from melanoma in adult fish into 2dpf larvae that were kept in embryo medium containing Aspirin or Celecoxib (COX-2 selective inhibitor). B) Representative images of melanoma invasiveness at 7dpi upon Aspirin treatment. C) Time-course of melanoma invasiveness in WT and G2 *tert−/−* larvae under Aspirin treatment. D) Slope of HRAS melanoma spread between 1, 4 and 7dpi. Comparison of invasiveness in a WT or G2 *tert−/−* either untreated (Control) or containing Aspirin (WT N=26 and G2 *tert−/−* N=29) or Celecoxib (WT N=19 and G2 *tert−/−* N=13). Each dot represents one zebrafish larva from 2-3 biological replicates.

## Discussion

Studies on how telomerase affects tumorigenesis have focused primarily on the cell-autonomous role of telomere shortening in cancer cells (9). Indeed, telomerase is reactivated in the majority of cancer cells promoting cancer development. Consistently, telomerase promoter mutations that result in increased telomerase expression are now recognized as one of the most common alterations in cancer (35). However, cancer incidence increases exponentially in the mid-ages of human life, a time when telomeres are shorter (1, 2). In our current study, we attempted to understand why cancer incidence increases when telomeres are shorter. Apart from the recognized cell autonomous role in tumor suppression, we propose that telomere shortening affects tumorigenesis in a non-cell autonomous manner. As an organism grows older, increasing numbers of cells with short telomeres modulate their surrounding environment creating a pro-inflammatory milieu that promotes tumorigenesis.

Using zebrafish embryo chimeras and cancer transplants, we show that incidence of melanoma is not only higher but progresses faster in animals deficient for telomerase. Both adult G1 *tert−/−* and G2 *tert−/−* embryos have shorter telomeres and mount DNA damage responses that stabilize p53 leading to premature aging and death (13, 28, 29). Indeed, mutations in *tp53* rescue the severity of both *tert−/−* models, allowing for prolonged survival. Spontaneous cancer in zebrafish, as in humans, is an age-associated disease that quickly rises upon decline of reproductive age (13, 36). Like other age-related phenotypes, spontaneous tumors in *tert−/−* zebrafish are accelerated to younger ages, while remaining similar in incidence and spectrum. Indeed, telomerase deficiency and telomere shortening in zebrafish do not appear to restrain tumorigenesis. Rather, they promote early cancer incidence denoting a systemic role in their effects. Similarly, humans with deficiencies in telomerase and premature telomere shortening show an increased cancer predisposition at younger ages (12). Thus, beyond preventing uncontrolled cell proliferation, absence of telomerase and telomere shortening appear to have a systemic role impairing health status and resistance to disease.

How could telomere shortening in surrounding tissues lead to increased incidence of cancer? We observed that *tert−/−* zebrafish present high levels of senescence. Studies *in vitro* revealed that senescent cells secrete SASP, composed by several inflammatory factors (14, 15). In agreement, we observed that *tert−/−* zebrafish present high levels of *cdkn1a* and *ink4ab* senescence genes and TNF-α, a cytokine involved in systemic inflammation. Moreover, G2 *tert−/−* cells are capable of inducing systemic senescence and inflammation in a non-cell autonomous manner. This data constitutes a strong indication that cells with short telomeres are a source of paracrine SASP *in vivo.* However, similar to other studies in zebrafish (37), we were unable to detect other typical SASP cytokines in *tert−/−* zebrafish larvae, such as IL6 and IL10. The main *in vivo* SASP molecules are yet to be identified in zebrafish.

Consistent with higher levels of inflammatory cytokines, G2 *tert−/−* cells containing critically short telomeres can modulate their environment by increasing the number of neutrophils. An increase in innate immune cells is characteristic of an inflammatory environment which can be tumorigenic. Human skin cancers have been shown to increase upon repeated injury and ulcers of previous lesions (38). In zebrafish, Feng *et al*. showed that preventing the recruitment of innate immune cells reduced the growth of HRAS^G12V^-transformed cells (16). Moreover, PGE2 produced by immune cells were shown to constitute a source of supportive signals for cancer cell growth. In line with this study, we observed a reduction of melanoma invasiveness with anti-inflammatory treatment, such as Aspirin and Celecoxib. Thus, our results suggest that G2 *tert−/−* cells with short telomeres promote the tumor invasiveness through the COX-2 pathway.

Collectively, our data indicates that an environment with short telomeres promotes tumorigenesis in a non-cell autonomous manner and increases the invasiveness capacity of melanoma cells. Apart from the recognized cell autonomous role in blocking uncontrolled cell division, telomere shortening and senescence may have a second, perhaps, antagonistic pleiotropic consequence of causing local tissue damage and chronic inflammation. Thus, we propose that telomere shortening during aging gives rise to a systemic inflammatory environment. Chronic inflammation may be part of the mechanism whereby telomere shortening leads to increase tumorigenesis with age. Indeed, whereas chronic inflammation was shown to be a contributing factor in several cancers, immunosuppression leads to increase the risk for certain tumors (39, 40). Epidemiology studies associate the long-term dosage of Aspirin with a reduced incidence of certain types of cancer (18–20). Interestingly, this effect is more pronounced with increased age of the population. Reverting telomere shortening in animal models that possess short telomeres, such as the zebrafish, will conclusively test the idea if repression of telomerase promotes cancer in aging.

## Materials and Methods

### Ethics statement

All Zebrafish work was conducted according to National Guidelines and approved by the Ethical Committee of the Instituto Gulbenkian de Ciência and the DGAV (Direcção Geral de Alimentação e Veterinária, Portuguese Veterinary Authority).

### Zebrafish maintenance and standard techniques

Zebrafish were maintained in accordance with Institutional and National animal care protocols. For normal line maintenance embryos were collected from crosses and kept in E3 embryo medium (5.0mM NaCl, 0.17mM KCl, 0.33mM CaCl, 0.33mM MgSO4, 0.05% methylene blue, pH 7.4) at 28ºC on a 14h light/10h dark cycle. At 5-6dpf larvae were transferred into a recirculating system at 28ºC, with a 14h light/10h dark cycle. For anesthesia, fish were immersed into tricaine methane sulfonate solution at 168μg/L (MS222 Sigma) and after the procedure placed back into system water. Their recovery was monitored until they regained normal swimming ability. Tricaine methane sulfonate was used at high concentration (200mg/L) to sacrifice fish. Larvae (until 7dpf) were sacrificed by placing them in ice-cold water for longer than 20min.

### Transgenic and mutant zebrafish lines

The telomerase mutant line *tert*^*AB/hu3430*^ generated by N-Ethyl-Nnitrosourea (ENU) mutagenesis (Utrecht University, Netherlands; Wienholds, 2004), has a T→A point-mutation in the *tert* gene and is available at the ZFIN repository, ZFIN ID: ZDB-GENO-100412-50, from the Zebrafish International Re-source Center – ZIRC. The *tert*^*hu3430*^ mutation was combined by genetic crossing in a *casper* background strain leading to the complete lack of pigmentation (41). For maintenance of this line, *casper; tert*^*AB/hu3430*^ was continuously outcrossed to *casper; tert*^*+/+*^. All recipient embryos used for the generation of HRAS chimeras, *tert*^*hu3430/hu3430*^ homozygous mutants (referred to as *tert−/−*) as well as their WT siblings were obtained by incrossing casper; *tert*^*AB/hu3430*^ animals. Donor embryos carry two different transgenes: Tg(*mitfa:HRAS*^*G12*^-*mitfa:GFP*; β*-actin:mGFP*)). They express GFP and a mutated and oncogenic version of human HRAS under a melanocyte-specific promoter *mitfa* causing strong hyperpigmentation and the formation of melanoma (23). We used a Tg(β*-actin:mGFP*) line with ubiquitous expression of membrane bound-GFP (mGFP) (42), since *mitfa:GFP* is only visible upon melanocyte development.

### Generation of zebrafish chimeras

Both donor and recipient embryos were manually dechorionated using forceps (not earlier than 16 cell-stage). Dechorionated embryos were maintained in transplant-media (14.97mM NaCl; 503μM KCl; 1.29mM CaCl2 · 2H2O; 150.63μM KH2PO4; 50μM Na2HPO4; 994.04μM MgSO4 ·7H2O) with penicillin/streptomycin (100U/ml penicillin and 100μg/ml streptomycine) in agarose-coated plates until 48hpf after which they were transferred into E3 embryo medium in non-coated petri dishes. Cell transfer from donor to recipient embryo was performed at blastula-stage using a hydraulic, manual microinjector (CellTram® vario, Eppendorf) with needles pulled from capillaries (TW100-4, World precision instruments, with a tip clipped off and polished of inner diameter at the tip 40-45μm) using a fluorescent stereoscope (Leica M205FA). Labelled donor cells (GFP+) were taken from Tg(*mitfa:HRAS^G12^-mitfa:GFP;* β*-actin:mGFP*) embryos and injected into recipient embryos. Cells were taken up by gentle suction directly at the blastula surface and released by injecting into the blastula of the recipient without ever harming the yolk cell. To increase the likelihood of transferring neural crest progenitors for tumor studies in adult animals, cells were typically taken from 3-5 spots at different sides of the donor embryo, all aligned midway between animal pole and yolk cell. Directly upon transfer, around 5% (estimation) of cells in a chimeric embryo were donor-derived. Single donor embryos served usually for various recipients (up to four), but one recipient never received cells from mixed donors. Upon cell transfer embryos were kept at low density (max 50 per plate) at 28ºC and cleaned daily.

### Selection of Chimeras to grow and tumor assessment

All animals included in this study were screened for a normal phenotype, presence of melanocytes and presence of GFP-positive donor cells. This screening was done under light anesthesia (84μg/L tricaine methane sulfonate MS222 in E3 embryo medium, 50% of standard concentration) under a Fluorescence stereomicroscope (Leica M205FA). Tumor appearance was assessed weekly and macroscopically. Individual animals were scored for the onset of a vertical growth phase, the presence of an outgrowth in any direction. Subsequently, most animals were analyzed by histopathology to confirm tumor formation and the state of invasiveness.

### Fish preservation for histology

When possible, fish were food-deprived for 24h prior to processing. After sacrificing, pictures of each fish were taken from both sides, both with a regular camera and at the fluorescent stereoscope (Leica M205FA) to save information about the gross distribution of pigmentation and chimeric (GFP+) cells. Animals were fixed in 10% neutral buffered formalin for 72h at room temperature and decalcified in 0.5M EDTA for 48h. Whole fish were paraffin embedded and 3μm transversal cuts were done from 5-8 regions of the fish (depending on size). Cuts were stained with haematoxylin and eosin and analyzed by histopathology. A total of N=18 animals was analyzed (9 WT and 9 *tert−/−* recipients).

### Melanoma cell transplants into 2dpf larvae

Melanoma cells were derived from Tg(*mitfa:HRAS*^*G12*^-*mitfa:GFP*) zebrafish tumors. To obtain the tumor cells fish were first sacrificed with tricaine 25x and the tumor was dissected with a regular scalpel and scissors. To dissociate the tumor, the mass of cells was dissected in small pieces, placed in a tryplE solution and pipetted up and down. Enzymatic reaction was stop with the addition of FBS (10% of total volume). Solution was filtered (70μm filter) and spun down at 1700 rpm for 5 min. The pellet was re-suspended in PBS calcium/magnesium free and then washed in culture medium with PBS (DMEM + 10% FBS). The final solution was approximately 1×10^7^ cells/mL and was obtained by removing as much as possible supernatant in the last centrifugation.

Melanoma cells were injected into the circulation of 2dpf larvae with a microinjection apparatus and needles were pulled from capillaries (TW100-4, World precision instruments). Transplanted larvae were kept overnight at 28ºC in embryo media. In the following day larvae are screened for the presence of GFP positive cells in the tail region and only those continue in the experiment. Pictures were taken at 1, 4 and 7 days-post injection using a fluorescent stereoscope (Leica M205FA). Transplanted larvae were kept in individual wells of a 6 well-plate to allow individual tracking of melanoma progression. Control (E3) or treatment (Aspirin - 30µM; Celecoxib – 25µM dissolved in DMSO) media were replaced daily. These experiments were repeated 2-3 times. GFP area was quantified using the Analyze Particles tool of imageJ 1.52i software.

### Statistical analysis

Statistical analysis was done with the Software GraphPad Prism 6. Comparisons of two different points were done by unpaired t-test. For the G1 HRAS chimeras, comparison over time (for at least 2 timepoints) was performed by Two-way RM ANOVA. Tumor onset over time was compared using a Log-rank (Mantel-Cox) test. A critical value for significance of p<0.05 was used throughout the study. For the larvae transplants, trend lines of GFP area between the three time-points (1, 4 and 7 days-post injection) per transplanted zebrafish were calculated using Microsoft Excel 2010 software. Slopes averages were compared between each two conditions using unpaired t-test with the Software GraphPad Prism 6.

### Senescence-associated β-galactosidase assay

β-galactosidase assay was performed as previously described (43). Briefly, sacrificed zebrafish larvae were fixed over-night in 4% paraformaldehyde in PBS at 4°C and then washed three times for 1 h in PBS-pH 7.4 and for a further 1 h in PBS-pH 6.0 at 4°C. β-galactosidase staining was performed for 10h at 37°C in 5 mM potassium ferrocyanide, 5 mM potassium ferricyanide, 2mM MgCl2 and 1 mg/ml X-gal, in PBS adjusted to pH 6.0. After staining, larvae were washed three times for 5 minutes in PBS pH 7, observed and photographed using a bright filter stereoscope (Leica M205FA).

### Real-time quantitative PCR

4dpf larvae were sacrificed, immediately snap-frozen in liquid nitrogen and collected in Eppendorf tube, minimum 10 larvae each. RNA extraction was performed using a RNeasy extraction kit (Qiagen, UK # 50974134). Briefly, larvae were smashed in RLT lysis buffer (provided by the kit) and the extract was washed and RNAs isolated through RNA binding column and eluted in dH_2_O RNase-free, according to manufacture procedures. Quality of RNA samples was assessed through BioAnalyzer (Agilent 2100, CA, USA). Retro-transcription into cDNA was performed using a RT-PCR kit NZY First-Strand cDNA Synthesis Kit # MB12501 (NZYtech). Quantitative PCR (qPCR) was performed using iTaq Universal SYBR Green Supermix # 1725125 (Bio-Rad) and an ABI-QuantStudio 384 Sequence Detection System (Applied Biosystems, CA, USA). qPCRs were carried out in triplicate for each cDNA sample. Relative mRNA expression was normalized to rpl13 mRNA expression using the DCT method. Primer sequences are listed in Table S1.

**Table S1.**
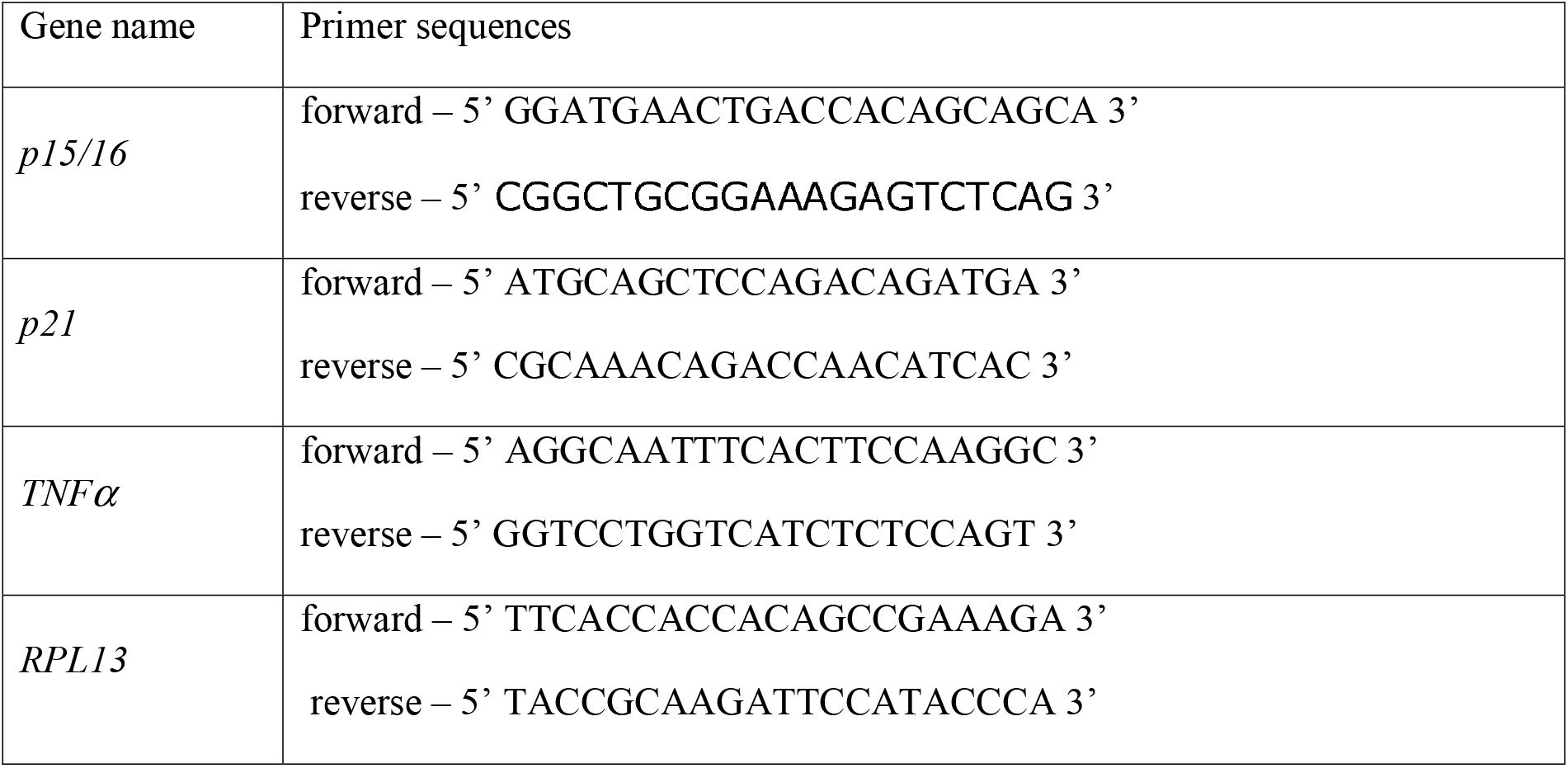
List of primers used in RT-qPCR expression analysis and *tert* genotyping.

## Acknowledgements

We thank members of the Telomeres and Genome Stability Laboratory for helpful discussions. We are grateful to Yi Feng (U of Edinburgh), Thiago Carvalho (Fundação Champalimaud) and Leonor Saúde (Instituto de Medicina Molecular) for critically reading our manuscript. We thank the Instituto Gulbenkian de Ciência histology unit and the Fish Facility for excellent animal care. KL was a recipient of a Portuguese Fundac□ão para a Cie□ncia e a Tecnologia (FCT) fellowship SFRH/BD/52173/2013. This work was supported by the FCT (PTDC/BIM-ONC/3402/2014 and PTDC/SAU-ONC/116821/2010) and the Howard Hughes Medical Institute grants received by MGF.

## Author contributions

Conceived and designed the experiments: MGF KL MF and MMG. Performed the experiments: KL MMG MF MM BLB KG. Analysed the data: KL MMG MF TC MGF. Contributed reagents/materials/analysis tools: KL MMG MF MM BLB KG TC MGF. Wrote the paper: MGF KL MMG.

---

**Supplementary Figure 1.**
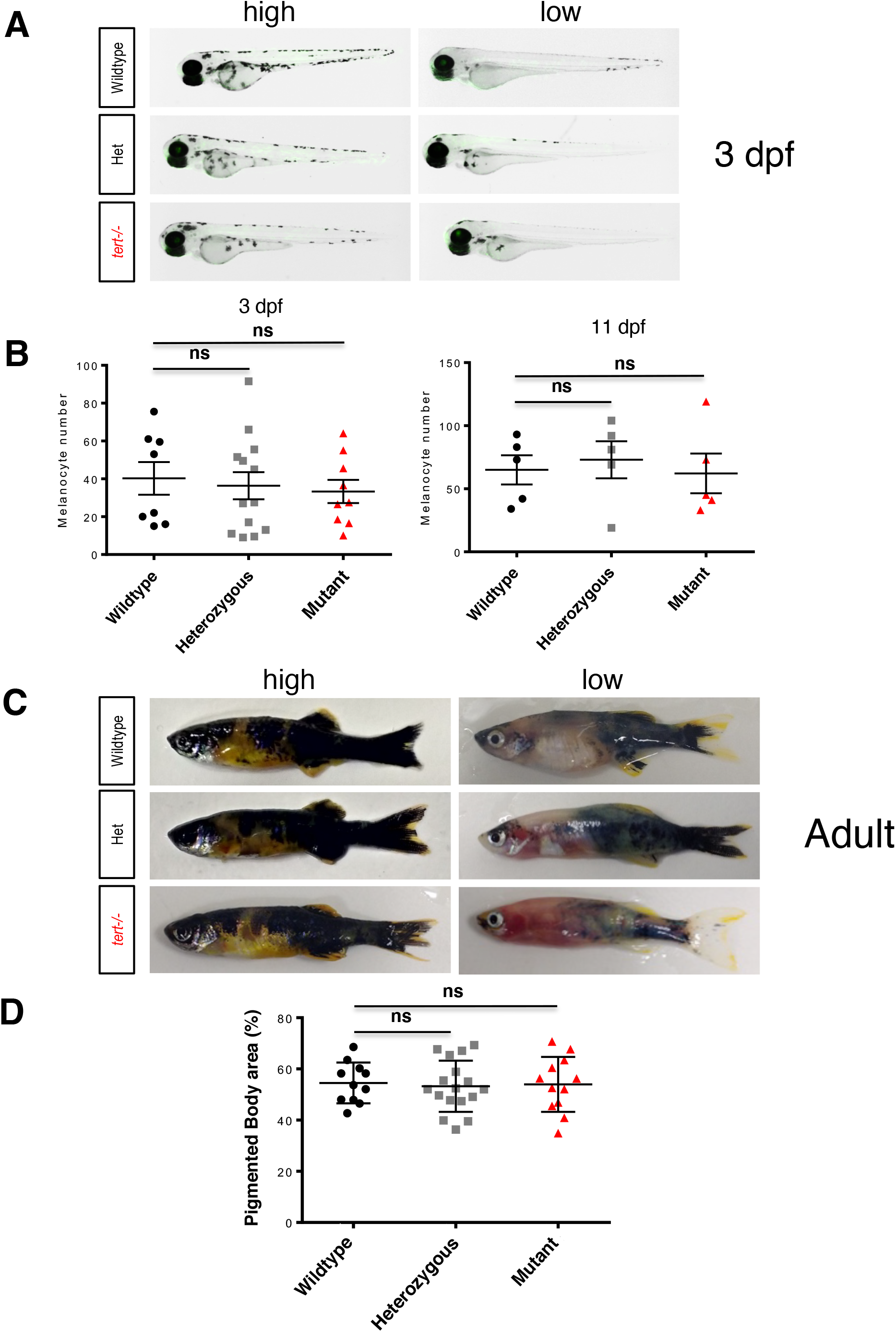
*tert* genetic status of chimera recipients does not influence the number of melanocytes in larvae or adults. A) Representative images of 3dpf chimeras exhibiting high (left) and low (right) number of melanocytes. Blastula mitfa:HRAS; β-actin:GFP cells were injected into the same stage embryos resulting from an incross of *tert+/−*; Casper. Larvae were genotyped at 3dpf and followed individually until 11dpf. B) No significant differences can be observed between the number of melanocytes in hosts of different *tert* genotype, both at 3dpf and 11dpf. Each point in the graph represents an individual animal. Data are represented as mean +/− SEM. C) Chimeras harboring a tumor were analyzed for extent of pigmentation in adults. Left side animals with high pigmentation, right side: low pigmentation. C) Quantification of pigmented area given as in percent of total surface. Each datapoint represents one animal (both sides). Pigmented area did not significantly differ depending on the host genotype. Data are represented as mean +/− SEM.

